# Meta-analysis: Purity of extracted human blood DNA, *E. coli* gDNA, and plasmid using magnetic nanoparticle

**DOI:** 10.1101/2024.10.15.618603

**Authors:** Stanley Evander Emeltan Tjoa, Mudasir Mudasir, Edi Suharyadi, Budi Setiadi Daryono

## Abstract

DNA extraction technology continues to evolve along with the need for a fast and simple process. The DNA extraction with magnetic nanoparticles accommodates these demands by providing a method that is safe, simple, and yet able to give good purity. The purity parameter of the extracted DNA is based on the A260/280 value. This meta-analysis aims to find variables that can affect the purity of extracted DNA using magnetic nanoparticles, based on its extraction and particle synthesis procedure. The information to be analyzed is from DNA extraction from human blood, plasmid extraction, and *E. coli* gDNA extraction. The research begins with a systematic review to collect appropriate studies and analysis will be carried out using Minitab to find variables that affect the purity of DNA, either positive or negative. Here we show, that no variables have been found significant to the purity of the extracted DNA in both the DNA extraction procedure steps and reagents. For the magnetic nanoparticle synthesis method, for extraction of *E. coli* gDNA, coating agent TEOS/TiO_2_; CoZnFe_3_O_4_ magnetic nanoparticles, and the interaction between CoZnFe_3_O_4_ and TEOS are found to be variables that influence the purity of the extracted DNA based on A260/280. Hopefully, this information can be used as a development idea, either for the magnetic nanoparticle synthesis procedure or the selection of reagents and DNA extraction procedures. In addition, the results of this analysis can also be used for consideration in an in-house magnetic nanoparticle-based DNA extraction assembly.

## 1. Introduction

DNA extraction is a fundamental procedure that supports molecular biology laboratory research. DNA is being used for various processes ranging from amplification [1], cloning [2], and identification [3]. These processes require a DNA extraction method. The DNA quality can be evaluated from its purity from the absorption ratio at wavelength 260/280. DNA itself has an absorption at 260 nm. Proteins that are commonly impurities have absorption at 280 nm. A value of 1.6 – 2 would indicate that an extract has good purity [4]. Purity is essential because many proteins in DNA extracts can interfere with subsequent processes.

A DNA extraction method that is currently being developed is the use of magnetic nanoparticles. A magnetic nanoparticle is designed in such a way that it can bind DNA. The magnetic particles will be bound by an external magnet. DNA bound to the magnetic modified particle can be separated from impurities remaining in the solution. In the final step, the DNA will be eluted with water or a suitable solution so that it can be separated from the magnetic nanoparticle. This technique is relatively simple compared to other DNA extraction methods. The use of magnetic nanoparticles for DNA extraction is widespread. The target samples used also varied. Starting from human blood [5-9] to bacterial samples, both genomic DNA (gDNA) [6,10-13] and plasmids [8, 14-22].

Research on magnetic nanoparticle-based DNA extraction has been widely carried out but is scattered. Researchers take various approaches to its development. This study aims to offer another approach for researchers that builds on existing studies in magnetic nanoparticle synthesis and DNA extraction methods.

In this research, we will systematically collect papers about magnetic nanoparticles synthesized for DNA extraction. We will categorize all the collected research and create groups based on the type of sample and target DNA. From there, we will look for any variables that influence the purity of an extracted DNA.

## 2. Methods

Literature collection uses keywords with the help of boolean operators and quotation marks: “magnetic nanoparticle” AND “DNA extraction” OR “DNA isolation” in four databases: JSTOR, DOAJ, Mendeley, and Scopus. The results are 1, 190, 10, and 166 articles, respectively. Next, screening was based on title, abstract, and full-text availability. There were 72 journals. Then eight titles were added using manual searches from Google Scholar.

In the next step, the contents of the journals were extracted. There were 45 titles eliminated on the grounds of using commercial nanoparticle magnets, DNA interaction studies using commercial DNA or commercial isolation kits, machine development, not having clear magnetic nanoparticle synthesis information, duplicate titles, and not showing A260/280 results.

In the final step, categorization was carried out based on the type of sample processed. There were five for blood samples, five for *E. coli* gDNA samples, and ten for bacterial plasmid extraction. Sixteen papers were omitted as they were not comparable due to differences in the target processed samples. We include the PRISMA statement in Figure 1 to illustrate the exclusion and inclusion stages of the articles used in this study.

**Figure 1.**
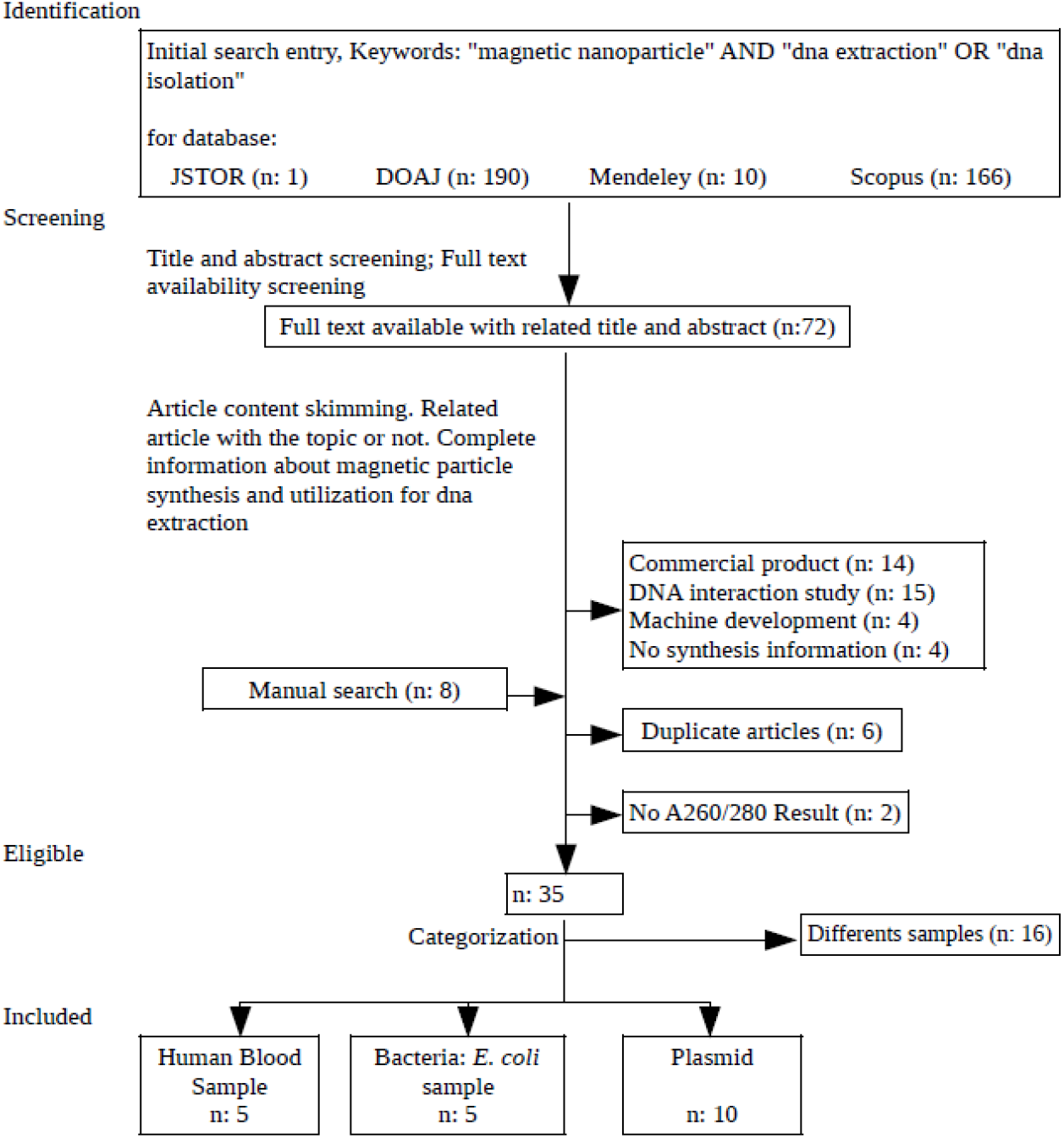
PRISMA Statement for Article Inclusion for This Study.

Data analysis was conducted using Minitab 19. All variables from each paper were explored and compared using Design of Experiment (DOE), factorial screening. The response is the A260/280 value. The data that would be obtained is a Pareto chart to determine the variables that have a significant effect and the main effect plot to see the pattern of the influence of each variable on the analyzed response.

## 3. Results and Discussion

Extraction using magnetic nanoparticles has been developed for a long time. Many commercial kits utilize magnetic nanoparticles for various types of samples [23]. The utilization of functional groups is also very diverse and ranges from proteins [20] to silica [6,11-13,15,21]. DNA extraction protocol, which covered the steps and reagents was analyzed. Pareto chart analysis was conducted for DNA extraction from human blood (Figure 2), *E. coli* gDNA extraction (Figure 3), and plasmid extraction (Figure 4), and none of the variables had a significant effect. Only the presence of guanidium HCl (GuHCl) almost crossed the reference line. GuHCl is a chaotropic salt that can form a salt bridge with a positive charge to enhance the interaction between silica and DNA which both have negative charges. The interaction is supported by low pH conditions, at ≤ 5. The interaction will decrease as the pH value increases [24].

**Figure 2.**
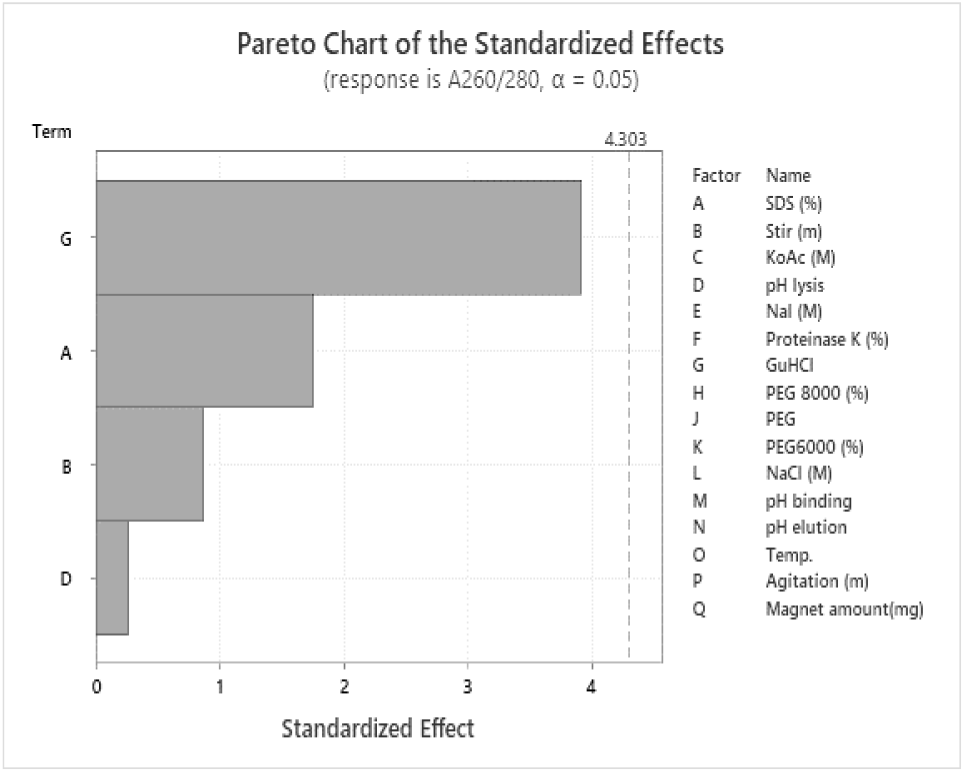
Pareto Chart of the Standardized Effects for DNA Extraction Protocol for Human Blood DNA.

**Figure 3.**
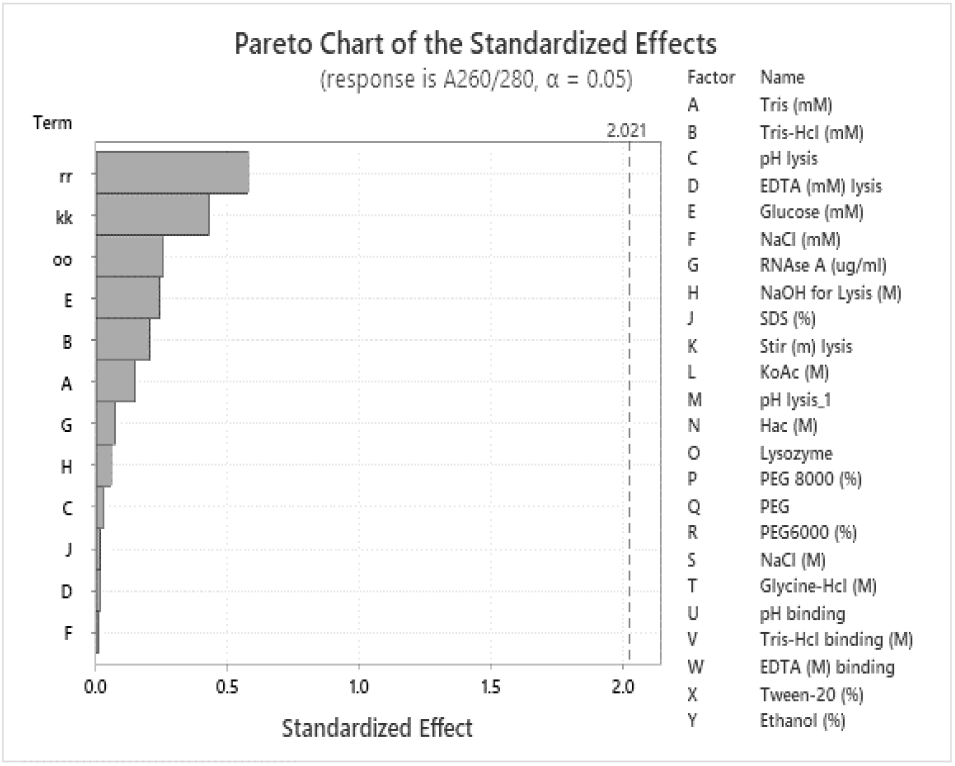
Pareto Chart of the Standardized Effects for DNA Extraction Protocol for *E. coli* gDNA.

**Figure 4.**
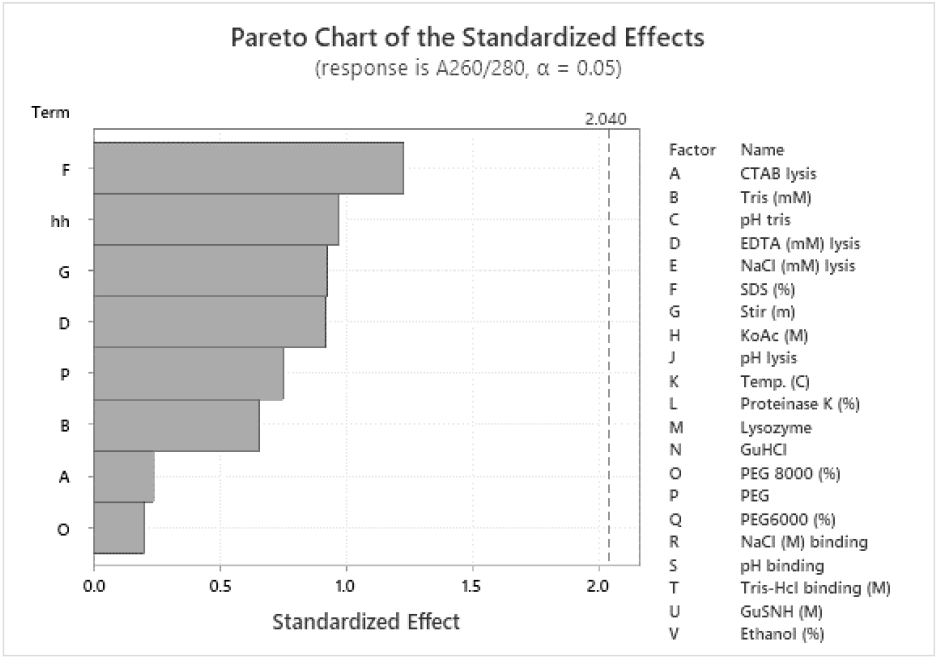
Pareto Chart of the Standardized Effects for DNA Extraction Protocol for Plasmid.

Main effect analysis was performed to obtain information about the negative and positive effects of each variable. DNA extraction in human blood samples (Figure 5), all variables were not significantly different. Although there is an increase and decrease, it is still within the range of 1.6 - 1.8. Only the GuHCl usage and the quantity of magnetic nanoparticles showed extreme changes. High levels of GuHCl caused a decrease in the purity ratio to below 1.5. GuHCl usage was consistent with the Pareto chart, while not significant, the presence of GuHCl affects the quality as viewed from the A260/280 response.

**Figure 5.**
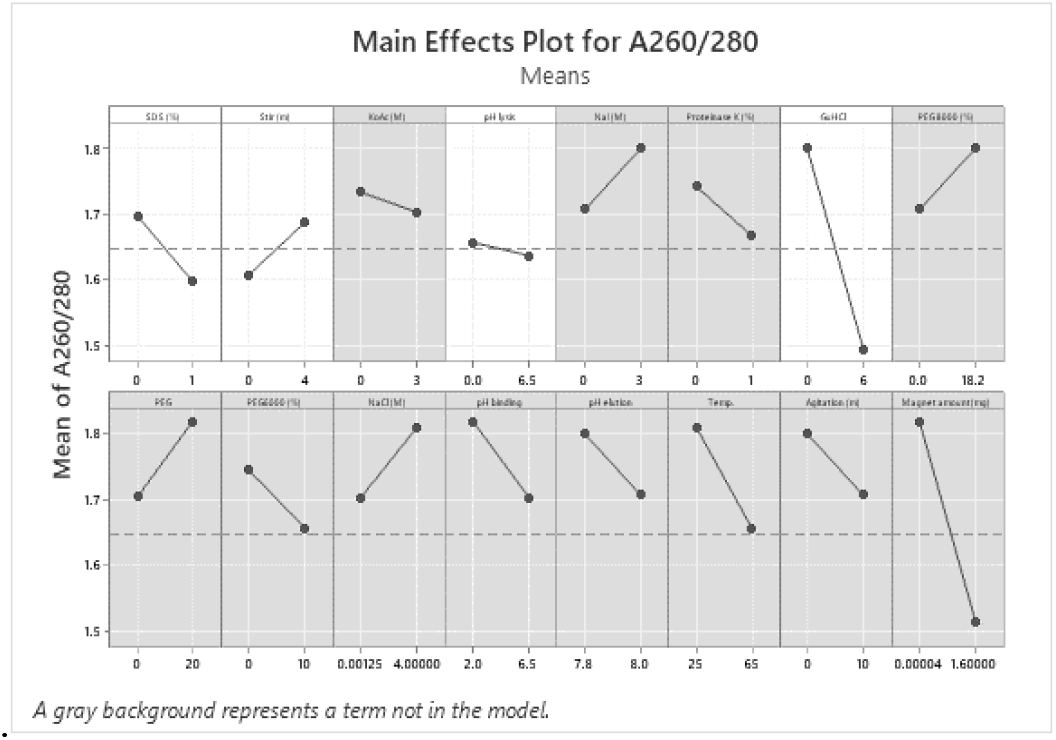
Main Effects Plot for A260/280 of DNA Extraction Protocol for Human Blood DNA.

As the level of GuHCl increased, the purity ratio of the DNA extract decreased. This could be caused by the phenomenon of positively charged salt bridge formation. Proteins have various charges. Thus, a positive charge salt bridge can cause the protein to bind to the silica surface. In addition, there are various other protein-silica interaction mechanisms [25-27]. The greater the magnetic nanoparticles used, the lower the A260/280 ratio. It could happen because the more magnetic nanoparticles, the more surface area interacts with the protein. So, the A260/280 value drops because the amount of protein in the extracted DNA increases.

The main effect analysis on plasmid extraction (Figure 6) provided information that the more EDTA was used in lysis, the lower the ratio value. The more, up to 2%, Sodium Dodecyl Sulfate (SDS) increased the A260/280 ratio value. In the *E. coli* gDNA extraction protocol (Figure 7), EDTA had a positive effect and the stir stage had a negative impact. The other variables showed no effect.

**Figure 6.**
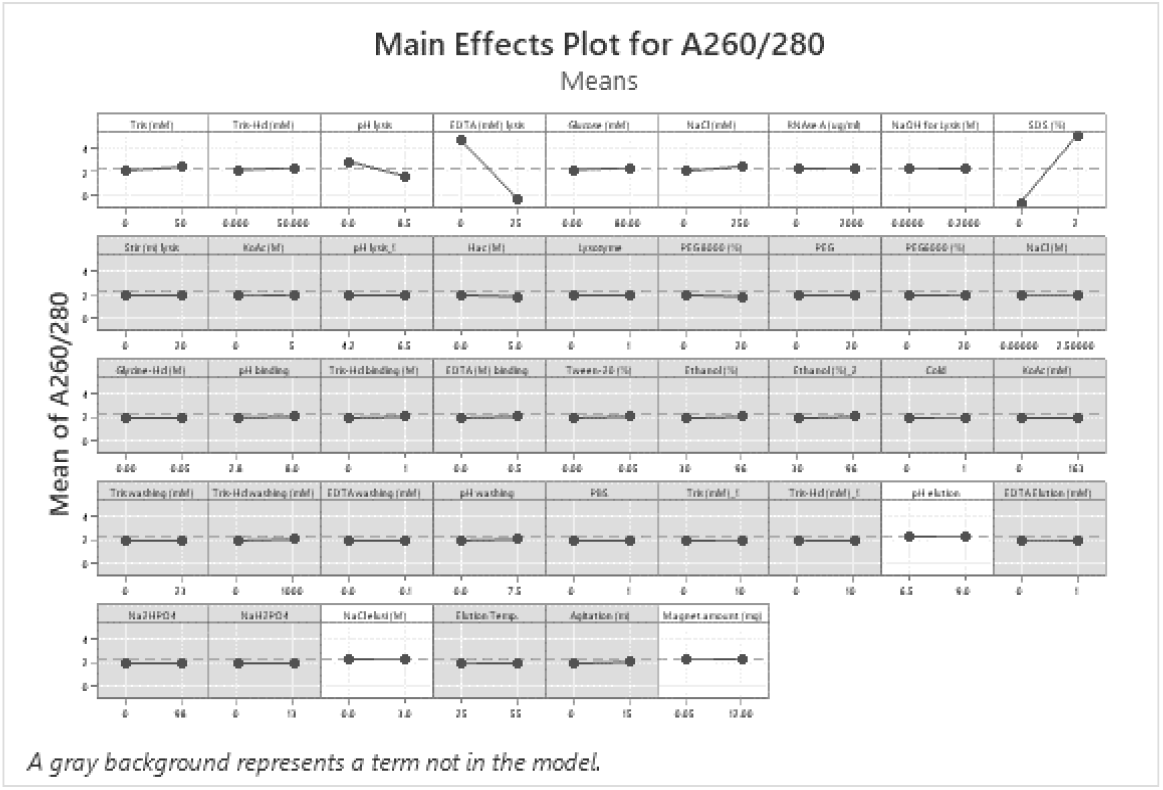
Main Effects Plot for A260/280 of DNA Extraction Protocol for Plasmid.

**Figure 7.**
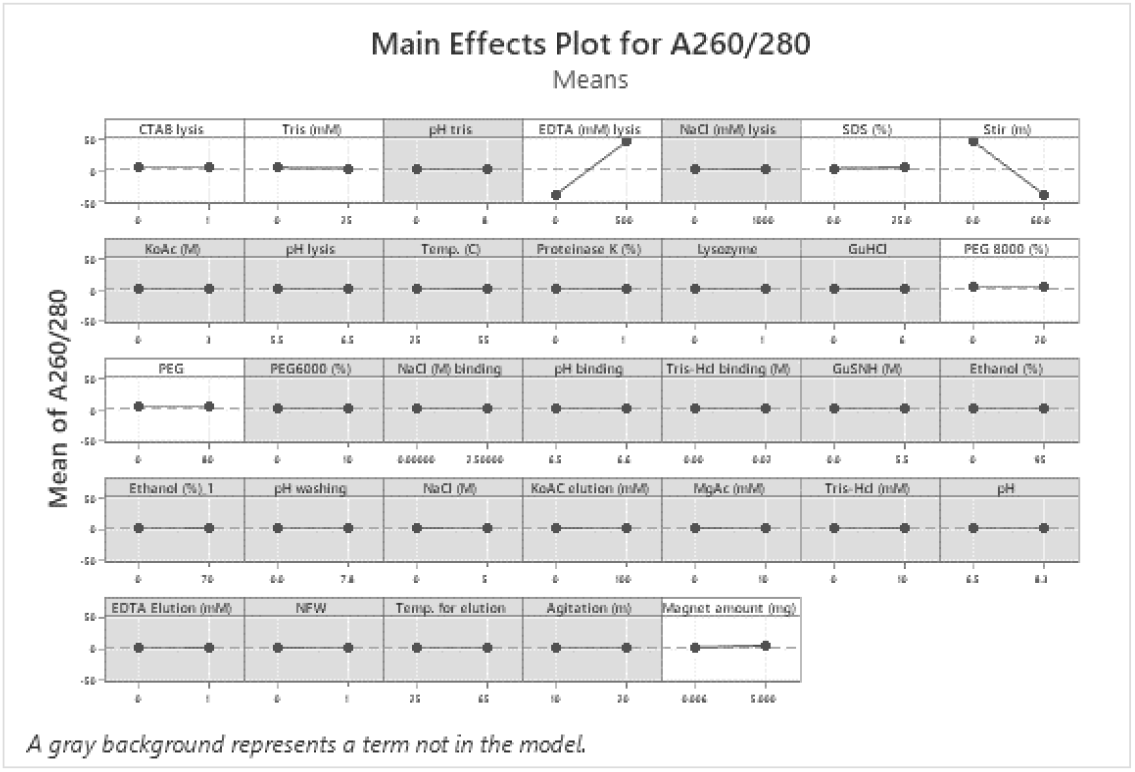
Main Effects Plot for A260/280 of DNA Extraction Protocol for *E. coli* gDNA.

EDTA in lysis buffer for plasmid and gDNA extraction has opposite effects on A260/280 values, decreasing and increasing respectively. Generally, EDTA functions as a chelator for cofactors in nuclease enzymes. Thus, it inhibits the activity of enzymes that degrade DNA. In addition, EDTA may cause precipitation of some proteins. Especially for proteins stabilized by metal ions, such as His-tag proteins. As a result, the proteins will denature, adsorb to the magnetic nanoparticles, and contaminate the extracted DNA. These two reasons may account for the different effects of EDTA on different sample types.

SDS functions as a lysing agent, to break down cells to allow DNA to be extracted. A small amount of SDS causes a small amount of extractable DNA and may also leave a lot of impurities in the form of proteins. This is also the case with the stirring time in the lysis step. The longer the stirring time, the more cells are broken, and consequently, the more impurities are degraded and can be separated.

The variables at the fabrication step of magnetic nanoparticles were also analyzed. In the Pareto chart, in the categories of whole blood DNA extraction (Figure 8), plasmid extraction (Figure 9), and *E. coli* gDNA extraction (Figure 10), TEOS/TiO_2_; CoZnFe_3_O_4_, and interaction between CoZnFe_3_O_4_ and TEOS gives a significant effect on *E. coli* gDNA extraction based on A260/280 value.

**Figure 8.**
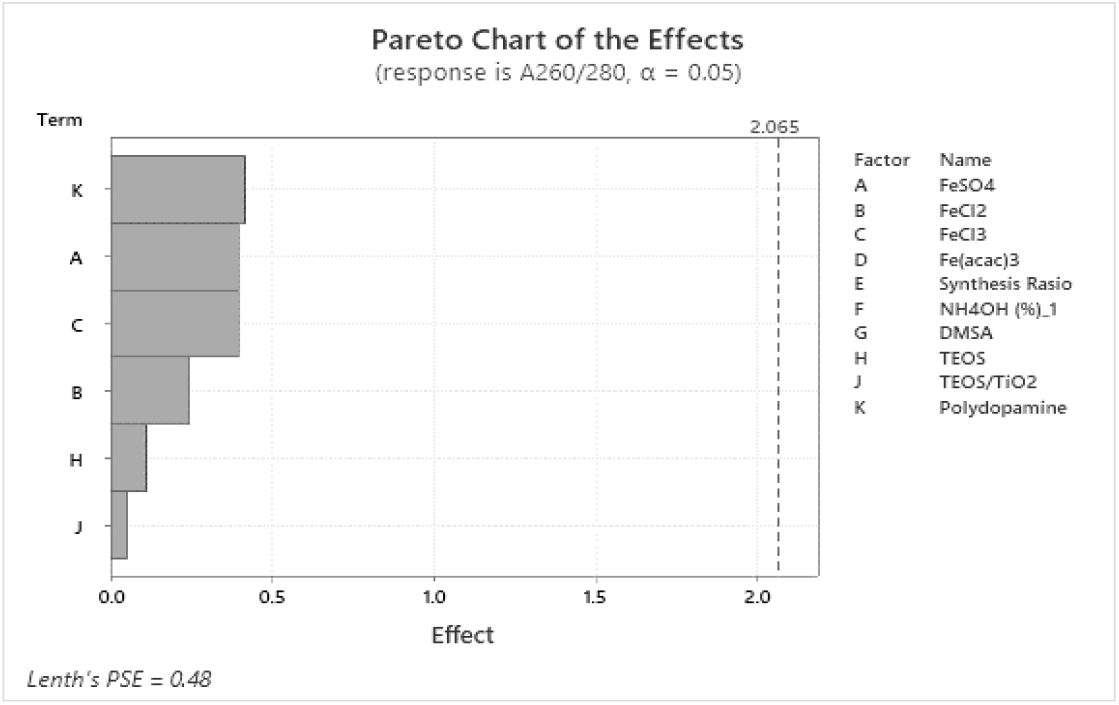
Pareto Chart of the Standardized Effects for Magnetic Nanoparticle Synthesis for Human Blood DNA Extraction.

**Figure 9.**
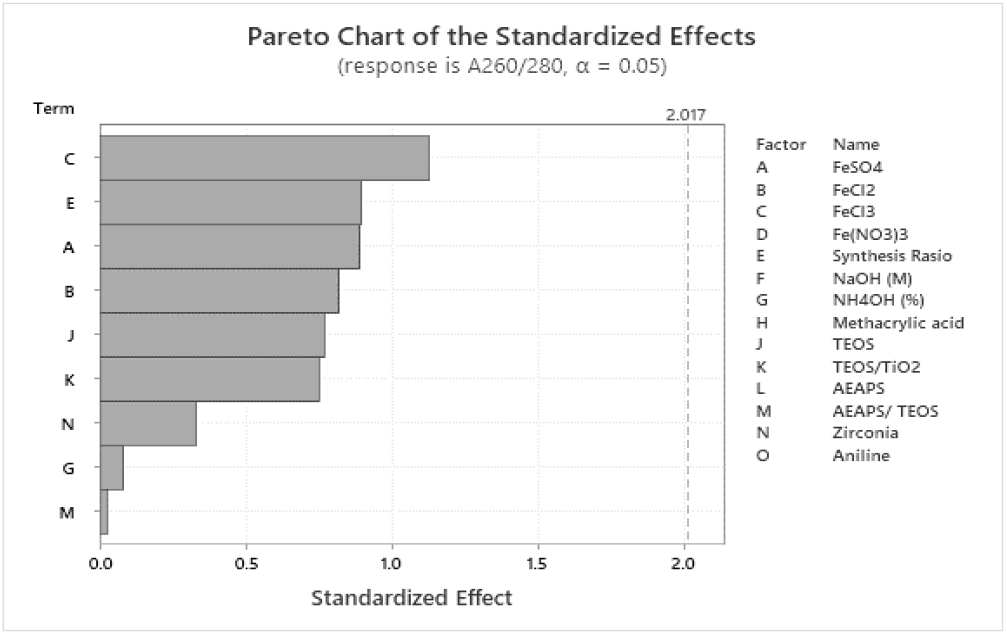
Pareto Chart of the Standardized Effects for Magnetic Nanoparticle Synthesis for Plasmid Extraction.

**Figure 10.**
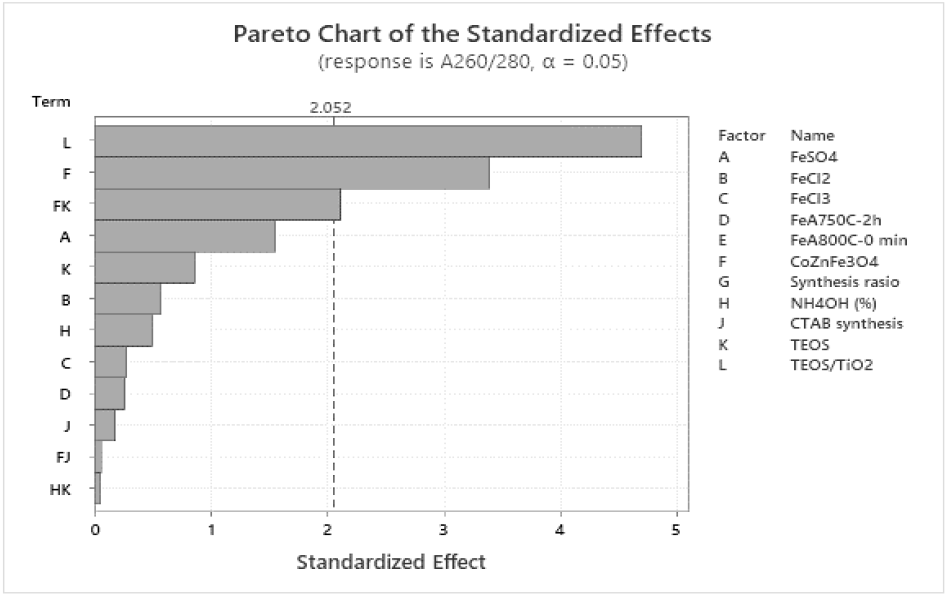
Pareto Chart of the Standardized Effects for Magnetic Nanoparticle Synthesis for *E. coli* gDNA Extraction.

In the main effect plot for the magnetic nanoparticle synthesis variable for human blood DNA extraction (Figure 11), the different Fe salts used affect the A260/280 values. The use of FeSO_4_ and FeCl_2_ had a positive effect and the use of FeCl_3_ had a negative result. Coating reagents, TEOS and TEOS/TiO_2_, and polydopamine showed positive effects. In the analysis for plasmid extraction (Figure 12), the results are similar to the investigation on DNA extraction in human blood, the type of Fe salt shows a shift in the A260/280 value. The use of FeSO_4_, FeCl_2_, and FeCl_3_ have a negative effect. The ratio of Fe in the synthesis has a positive outcome. Other variables did not influence as seen from the range of A260/280 results ranging from 1.6 - 2. Analysis for *E. coli* gDNA extraction showed a positive outcome of Fe salt type. The use of TEOS/TiO_2_ coating agents has a negative result. The plot can be seen in Figure 13.

**Figure 11.**
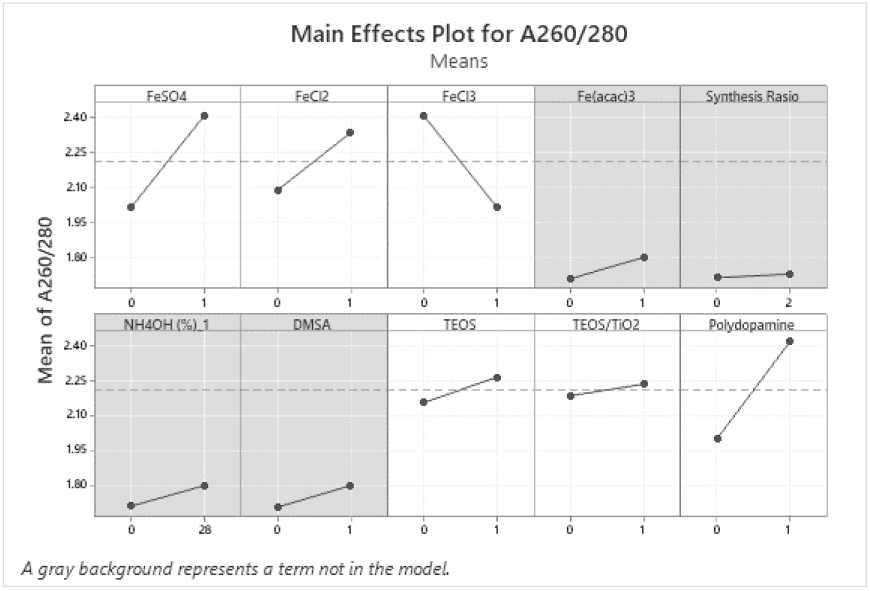
Main Effects Plot for A260/280 of Magnetic Nanoparticle Synthesis for Human Blood DNA Extraction.

**Figure 12.**
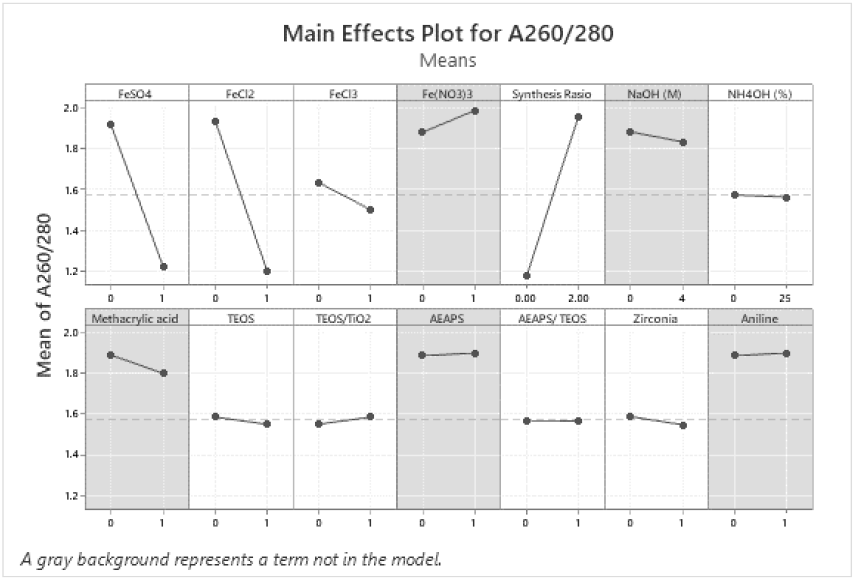
Main Effects Plot for A260/280 of Magnetic Nanoparticle Synthesis for Plasmid Extraction.

**Figure 13.**
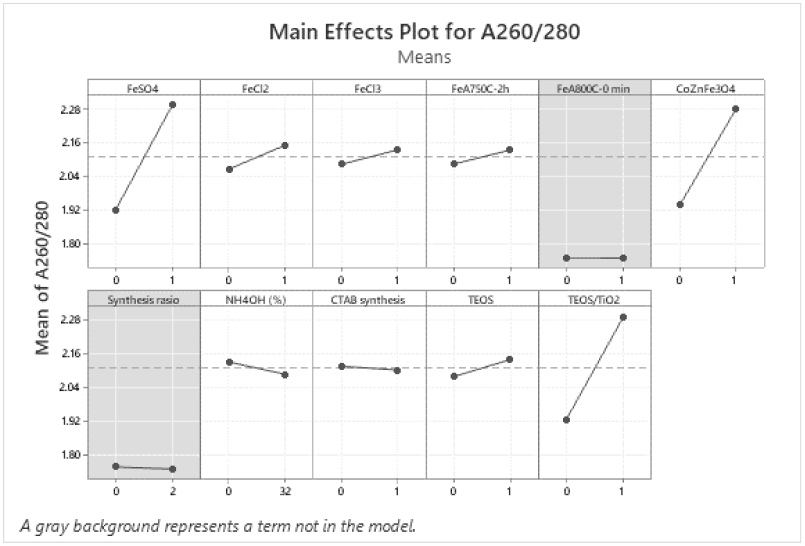
Main Effects Plot for A260/280 of Magnetic Nanoparticle Synthesis for *E. coli* gDNA Extraction.

Titanium oxide (TiO_2_) can bind to DNA strongly and spontaneously. Therefore, for DNA extraction, TiO_2_ is a good option as a coating agent [28]. In addition, TiO_2_ can also bind amino acid groups such as proline, lysine, and leucine [29]. Hal ini akan mempengaruhi kemurnian extracted DNA berdasarkan A260/280. If DNA purity is the main focus rather than yield, perhaps the use of TiO_2_ as a coating agent is not a preferred one.

CoZnFe_3_O_4_ can be used for magnetic nanoparticle precursors because it is not easily oxidized compared to Fe_3_O_4_ [12]. However, when viewed from the main effect plot, the use of CoZnFe_3_O_4_ results in a higher A260/280 value. The effect is also influenced by its interaction with tetraorthosilicate (TEOS). The silica surface produced by reaction with TEOS is known to affect its interaction with DNA/protein according to the binding buffer conditions [24-27].

In the synthesis of magnetic nanoparticles, there are several parameters that affect the shape and size of the particles. These range from the type of Fe salt to its ratio [30]. It could be that these different properties influence the amount of DNA/protein that can interact with the magnetic particles and thus affect the purity of the DNA extract. Although these variables have an insignificant effect.

Overall, it is known that the quality of extracted DNA based on A260/280 is more affected by magnetic nanoparticle synthesis variables. Other variables in the extraction protocol may play a role but are not significant. The application of magnetic nanoparticles to other samples can be initiated by considering the characteristics of the sample to be extracted. We hope these findings will help researchers develop protocols for utilizing magnetic nanoparticles in DNA extraction.

## 4. Conclusions

The application of magnetic nanoparticles in DNA extraction from blood samples, plasmid extraction, and E. coli gDNA extraction using standard protocols is quite robust. So far, no variables were found to be significant to the purity of the extracts. For the magnetic nanoparticle synthesis method, for E. coli gDNA extraction, the coating agent TEOS/TiO_2_, magnetic nanoparticles CoZnFe3O4, and the interaction between CoZnFe_3_O_4_ and TEOS were found to be variables that affect the purity of extracted DNA based on A260/280. The information obtained from this study can be supplementary to the new findings from recent references, especially since topics such as magnetic nanoparticle synthesis for DNA extraction are rapidly evolving.

## Acknowledgement

We would like to thank PT. Konimex for the support.

